# Chemical control of a CRISPR-Cas9 acetyltransferase

**DOI:** 10.1101/176875

**Authors:** Jonathan H. Shrimp, Carissa Grose, Stephanie R. T. Widmeyer, Ajit Jadhav, Jordan L. Meier

## Abstract

Lysine acetyltransferases (KATs) play a critical role in the regulation of transcription and other genomic functions. However, a persistent challenge is the development of assays capable of defining KAT activity directly in living cells. Towards this goal, here we report the application of a previously reported dCas9-p300 fusion as a transcriptional reporter of KAT activity. First we benchmark the activity of dCas9-p300 relative to other dCas9-based transcriptional activators, and demonstrate its compatibility with second generation short guide RNA architectures. Next, we repurpose this technology to rapidly identify small molecule inhibitors of acetylation-dependent gene expression. These studies validate a recently reported p300 inhibitor chemotype, and reveal a role for p300’s bromodomain in dCas9-p300-mediated transcriptional activation. Comparison with other CRISPR-Cas9 transcriptional activators highlights the inherent ligand tuneable nature of dCas9-p300 fusions, suggesting new opportunities for orthogonal gene expression control. Overall, our studies highlight dCas9-p300 as a powerful tool for studying gene expression mechanisms in which acetylation plays a causal role, and provide a foundation for future applications requiring spatiotemporal control over acetylation at specific genomic loci.

## Introduction

Lysine acetyltransferases (KATs) catalyze protein acetylation, a reversible posttranslational modification (PTM) that plays a critical role in many processes, including gene expression.^1^ Two of the most well-studied KATs are EP300 and its homolog CREBBP (commonly referred to jointly as p300/CBP). These two KATs possess a versatile substrate scope which includes histones, transcription factors, and members of the transcriptional regulatory apparatus itself.^2^ Accordingly, disruption of p300/CBP is associated with substantial changes in gene expression, and has been linked to several diseases.^3–4^ Besides its KAT domain, p300 and CBP additionally contain several non-catalytic modules including zinc fingers, acetylysine readers (bromodomain, BRD), methyllysine readers (PHD domain), and protein-protein interaction domains.^2^ Thus, a significant challenge in the study of p300/CBP lies in defining the specific role of the KAT domain in gene expression, as well as its targetable role in disease.

Considering methods to study cellular KAT activity, we were inspired by a recent report by Gersbach et al. which found that p300 could be delivered to specific genomic loci using the genomic-targeting methodology CRISPR-Cas9.^5^ Specifically, this study engineered a catalytically inactive variant of *S. pyogenes* Cas9 (dCas9) fused to truncated p300 module containing the BRD and KAT domains (dCas9-300) (Figure 1). Expression of this fusion in combination with chimeric short guide RNAs (sgRNAs) targeted to promoter regions led to induction of several model genes. Critically, control studies found inducible gene expression to be absolutely dependent on p300 KAT activity, as it was abolished by KAT-inactivating mutations.^5^ In addition to expanding the CRISPR gene activation toolbox, a broader implication of this study was that KAT activity can play a causative role in activation of gene expression, an often accepted premise for which limited unambiguous evidence exists.^6–7^ Moreover, these studies suggested dCas9-p300 may represent a valuable tool for studying this phenomenon in cells.

**Figure 1.**
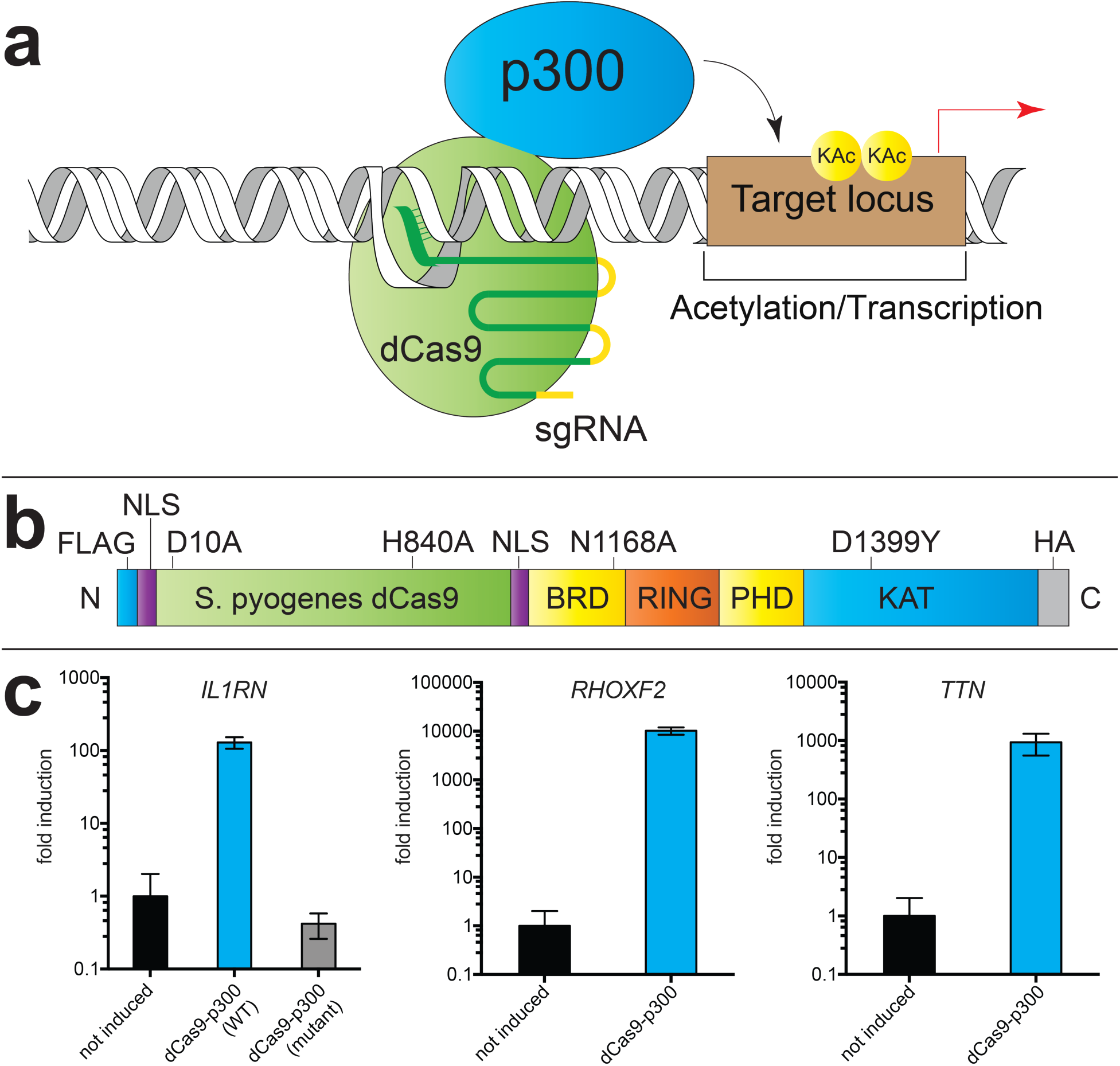
A CRISPR-Cas9 transcriptional reporter of KAT activity. (a) Scheme for induction of acetylation/gene expression by dCas9-p300, first reported by Gersbach et al. (b) Domain architecture of dCas9-p300 fusion. (c) Activation of model inducible genes *IL1RN*, *RHOXF2*, and *TTN* by dCas9-p300 in HEK-293 cells.

Here we describe the application of dCas9-p300 as a cellular reporter of KAT-dependent transcription. Building on previous studies, we first recapitulate dCas9-p300-mediated transcriptional activation, and benchmark its activity relative to non-KAT dCas9 gene expression platforms. Next, we demonstrate p300’s transactivation potential can be targeted to a model gene (*IL1RN*) in a dCas9-dependent manner using diverse sgRNA and fusion protein architectures. Finally, we apply this platform to analyze small molecules for disruption of KAT-dependent gene expression, highlighting the unique ligandability of dCas9-p300 relative to other transcriptional activators. Overall, our studies showcase the utility of dCas9-p300 as a tool for studying KAT-dependent transcription, and provide a foundation for its use in screening and epigenome editing applications.

## Results and Discussion

To apply dCas9-p300 as a transcriptional reporter of KAT activity, we first assessed its utility in cellular experiments. Consistent with previous reports,^5^ transient overexpression of dCas9-p300 in combination with sgRNAs tiling the *IL1RN* promoter strongly activated expression of this model gene in HEK-293T cells without modifying bulk acetylation (Figures 1, S1). Induced transcription of *IL1RN* was time-dependent, and continued to increase up to 72 h post-transfection (Figure S1). We found several other genes (*RHOXF2*, *TTN*) could also be induced by dCas9-p300, albeit to varying extents (Figure 1c). No induction of *IL1RN* was observed when an identical experiment was performed using dCas9-p300 D1399Y, a construct in which a key aspartate involved in the binding of acetyl-CoA to the p300 catalytic domain is mutated (Figure 1c). Of note, analysis of genomics data reveals that D1399Y is the most commonly observed missense mutations in p300 in human cancer, indicative of its ability to disrupt KAT-dependent p300 tumor suppression.^8^ This demonstrates the ability of dCas9-p300 induced transcription of *IL1RN* to be applied as a cellular reporter of KAT activity.

Next we sought to benchmark dCas9-p300’s transcriptional activation relative to other recently reported dCas9-based methods.^9–12^ The goal of these studies was two-fold: (i) to directly compare genomic recruitment of enzyme catalysis (dCas9-p300) versus recruitment of protein-protein interaction motif (dCas9-VP64) as transactivation mechanisms, and (ii) to obtain insights into the ability of dCas9-p300 to utilize alternative sgRNA architectures, an area that has not been previously explored. To accomplish this, we compared the ability of dCas9-p300 to activate *IL1RN* relative to components of the “Synergistic Activation Mediator” (SAM) platform, a method that has been powerfully applied in genome-wide screens.^10^ SAM-mediated gene expression is based on co-targeting of dCas9-VP64 and two MS2-HSF-p65 fusion proteins to genomic loci via an MS2 aptamer-containing short guide RNA (referred to as a sgRNA 2.0; Figure 2/S2). To compare these two approaches, we transiently transfected HEK-293 cells with components of the SAM platform or dCas9-p300 using sgRNA 2.0’s targeting the highly inducible *IL1RN* promoter. The strength of *IL1RN* induction followed the order SAM ≥ dCas9-p300 >> dCas9-VP64 (Figure S2). This rank order is consistent with previous reports, although these studies did not compare SAM and dCas9-p300 in the context of the highly inducible *IL1RN* locus.^13^ Interestingly, we did not observe a large effect upon replacement of dCas9-VP64 with dCas9 using the SAM platform, suggesting recruitment of MS2-HSF-p65 is sufficient to induce transcription when targeted to the *IL1RN* promoter. Importantly, dCas9-p300 promoted strong expression in these experiments, indicating its compatibility with second generation short guide RNA architectures.

**Figure 2.**
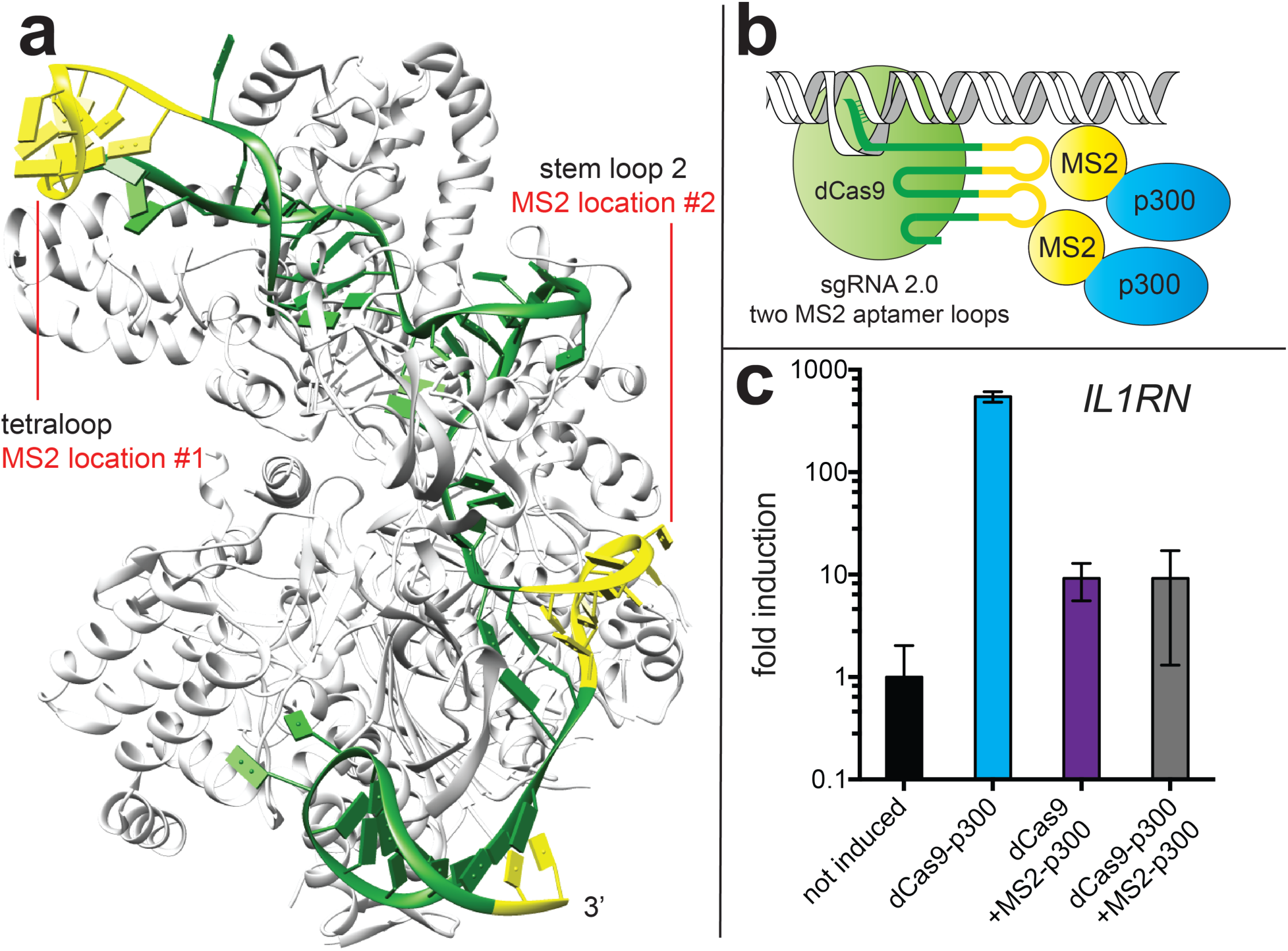
KAT-dependent gene expression can be stimulated using diverse genomic recruitment strategies (a) Structure of Cas9-sgRNA complex illustrating sites for short guide RNA engineering. (b) Cartoon schematic of dCas9-dependent recruitment of MS2-p300 fusions using sgRNA 2.0. (c) Comparison of KAT-dependent expression of *IL1RN* using dCas9-p300 and MS2-p300 with sgRNA 2.0 architectures.

Motivated by this finding, we next explored the ability of dCas9-sgRNA 2.0 complexes to mediate multivalent recruitment of p300 KATs to the *IL1RN* locus (Figure 2). In contrast to dCas9, which binds to genomic loci as a monomer, MS2 coat proteins bind their cognate aptamer as dimers,^14^ allowing a single sgRNA 2.0 to potentially recruit four copies of p300 catalytic activity. Thus, we designed a construct in which a CMV promoter drives constitutive expression of an MS2 coat protein fused at its C-terminus to the p300 core (Figure 2b). The p300 core used in these studies consists of the BRD, RING, and KAT domains, and is identical to that found in the dCas9-p300 construct. Employing this construct in combination with dCas9 and sgRNA 2.0s targeting the *IL1RN* promoter resulted in modest induction of gene expression after 72 hrs (Figure 2c). The less efficient transactivation of gene expression by MS2-p300 suggests it may be less efficiently recruited, less active, or improperly oriented at the *IL1RN* promoter relative dCas9-p300. Of note, recent studies employing a dCas9-Suntag construct to recruit the methylcytidine dioxygenase TET1 found the spacing of dCas9 recruited proteins to be essential for efficient recruitment of TET1 catalysis at specific genomic loci,^15^ suggesting a rationale for inefficient transactivation by MS2-p300. Indeed, in contrast to the SAM system, we found that coincident expression and recruitment of dCas9-p300 and MS2-p300 fusions squelched the induction of gene expression promoted by either component alone, consistent with counterproductive effects (Figure 2c). These studies provide insights into the ability of alternative sgRNA and fusion protein constructs to target KAT activity to specific genomic loci.

Next, we sought to apply dCas9-p300 as a tool to define the effect of small molecules on KAT-dependent transcription (Figure 3). We envisioned such an application of dCas9-p300 could fulfill two purposes. First, molecules directly inhibiting p300 KAT domain would be expected to decrease *IL1RN* induction, providing a cellular readout of p300 activity and rapid assessment p300 target occupancy, which has been difficult to assess using other approaches.^16–17^ Second, this approach should be amenable to the identification of molecules interacting with the transcriptional regulatory apparatus downstream of dCas9-p300, and thus have the potential to illuminate biological processes specifically necessary for KAT-dependent transcription. To explore this strategy, we screened a panel of small molecules for modulation of dCas9-p300 induced gene expression (Figure 3). Included in this initial panel were molecules interacting directly with acetylation-dependent signaling such as KAT, bromodomain, and histone deacetylase (HDAC) inhibitors, as well as ligands we hypothesized may target downstream biological processes necessary for dCas9-p300 transactivation, such as RNA Polymerase II (RNA Pol II) C-terminal domain phosphorylation and Polycomb Repressive Complex methyltransferase activity.^18–21^ To maximize dynamic range, treatments were carried out 48-72 h post-transfection, over which time *IL1RN*-induced gene expression increased ∼10-fold (Figure S1/S3). Using this approach, we observed the majority of molecules tested had only modest effects on dCas9-p300 induction of *IL1RN*. This may stem partly from the choice of concentrations used in our single point screen, as we purposely assessed inhibition of dCas9-p300 at concentrations determined to be non-toxic in HEK-293T in order to minimize off-target effects (Figure S3b). Three molecules in the panel demonstrated the strongest inhibitory effects: NCGC00496795, (+)-JQ1 (JQ1), and SGC-CBP30 (CBP30).^19–20, 22^ NCGC00496795 is a racemic member of a family of spirocyclic p300 inhibitors that were recently reported in the patent literature.^22^ Follow-up studies validated its ability to inhibit both p300 biochemical activity as well as dCas9-p300 driven gene expression in a dose-dependent fashion (Figure S4). Although we defer a full characterization of this compound class to future publications,^23^ the potent inhibition of dCas9-p300-induced *IL1RN* expression are consistent with the reported ability of this chemotype to target KAT domains, and validate our assay’s ability to identify p300 inhibitors.

**Figure 3.**
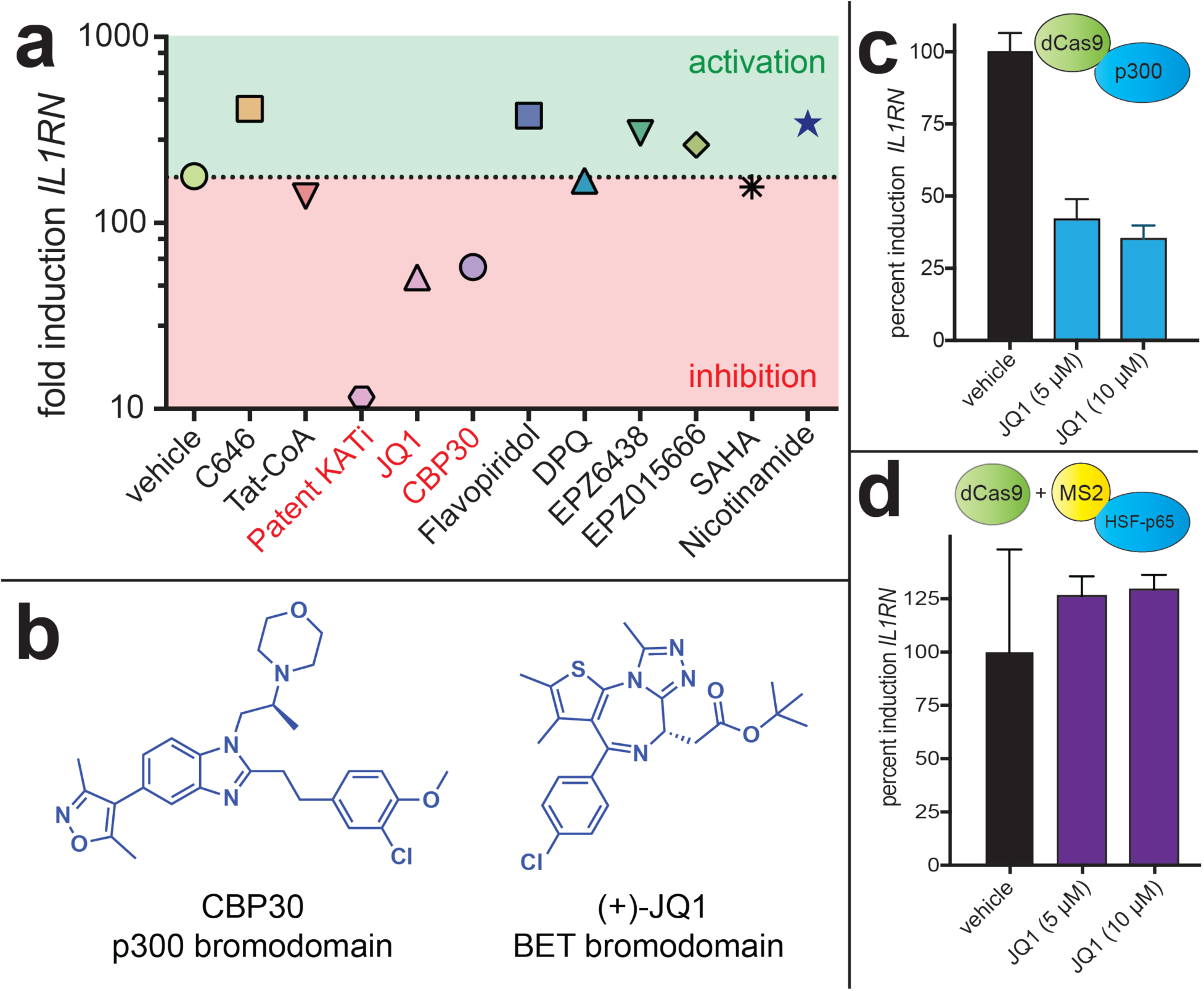
Influence of small molecule stimuli on CRISPR-Cas9 acetyltransferase. (a) Single point screening data. Single point concentrations are provided in the Supporting Information. (b) Structures of bromodomain-targeting small molecules identified as inhibitors of dCas9-p300-dependent gene expression. (c) Dose-dependent inhibition of KAT-dependent gene expression by JQ1. (d) JQ1 alters dCas9-p300-dependent gene expression but not SAM-dependent gene expression.

Notably, the other two molecules identified as dCas9-p300 inhibitors target bromodomains, small protein motifs that specifically bind acetylated lysine residues (Figure 3b).^24^ JQ1 is a chemical probe of the BET bromodomain family, which includes BRD2, BRD3, BRD4, and BRDT.^19^ Acetylation-dependent recruitment of BRD4 to genes is associated with RNA Pol II phosphorylation and transcript elongation, common features of activated gene expression. To build on the results of our pilot screen, we first confirmed dose-dependent inhibition of dCas9-p300-mediated gene expression by JQ1 (Figure 3c). We observed JQ1 concentrations as low as 5 μM were sufficient to inhibit *IL1RN* expression (∼30%). Controls verified this was not due to inhibiton of dCas9-p300 expression, which we found to be slightly upregulated by JQ1 treatment (Figure S5). Next, we performed experiments to assess whether BET bromodomain binding constitutes a specific or general requirement for dCas9-mediated transactivation. To accomplish this, we compared JQ1’s ability to inhibit *IL1RN* gene expression driven by either i) dCas9-p300 or ii) components of the SAM system (dCas9/MS2-HSF-p65). Interestingly, while JQ1 antagonized dCas9-p300, it had no effect on SAM-driven *IL1RN* expression (Figure 3d). This is consistent with the view that distinct transcriptional activation domains may exhibit unique cofactor dependencies.^25–26^ These studies indicate dCas9-p300 is uniquely dependent on acetylation-dependent protein-protein interactions relative to other dCas9-based transcriptional activators.

Next we turned our attention to understanding the effects of CBP30, a recently reported selective probe of the p300/CBP bromodomain.^20^ CBP30 binds the p300/CBP bromodomain with ∼40-fold selectivity relative to BET family members and has demonstrated phenotypic effects on p53 and IRF4-dependent transcription in cells.^20, 27^ Although it also targets an acetylation-dependent protein-protein interaction, CBP30 is unique relative to JQ1 in that it has the potential to directly interact with the BRD-containing dCas9-p300 construct. Again, we initially validated the effects of CBP30 on dCas9-p300 driven gene expression. CBP30 demonstrated dose-dependent inhibition of *Il1RN* transcription at concentrations similar to JQ1 (Figure 4a). Western blot analysis indicated this inhibition was not due to changes in ectopic expression of dCas9-p300 (Figure 4b). The structurally distinct p300/CBP bromodomain inhibitor SGC-CBP112 (CBP112) exhibited similar inhibition (Figure 4a).^28^ One notable feature of these studies was that relatively high concentrations of CBP30/CBP112 were required to inhibit *IL1RN* induction. This may reflect the fact that the target of these molecules (p300 BRD) is being constitutively overexpressed, limiting their potency, or alternatively indicate engagement of an off-target. Therefore, to more conclusively examine the role of the BRD in dCas9-p300-mediated gene expression we prepared a construct harboring a bromodomain mutation (CBP, N1168A; p300 N1132A) known to abrogate acetyllysine binding (Figure 4c).^29^ Consistent with the effects of CBP30, the dCas9-p300 bromodomain mutant showed diminished transactivation potential (Figure 4d-e). Of note, previous studies have suggested the bromodomain of p300 may alter the in cis catalytic activity of the acetyltransferase domain,^30–31^ providing a rationale for the observed effects. However, it is important to note that our studies do not definitively rule other contributions of the p300 bromodomain to dCas9-p300 mediated transcription, including effects on processive acetylation or genome-wide localization.^32^ These studies establish a role for the bromodomain in fine-tuning KAT-dependent transcriptional activation at the *IL1RN* locus.

**Figure 4.**
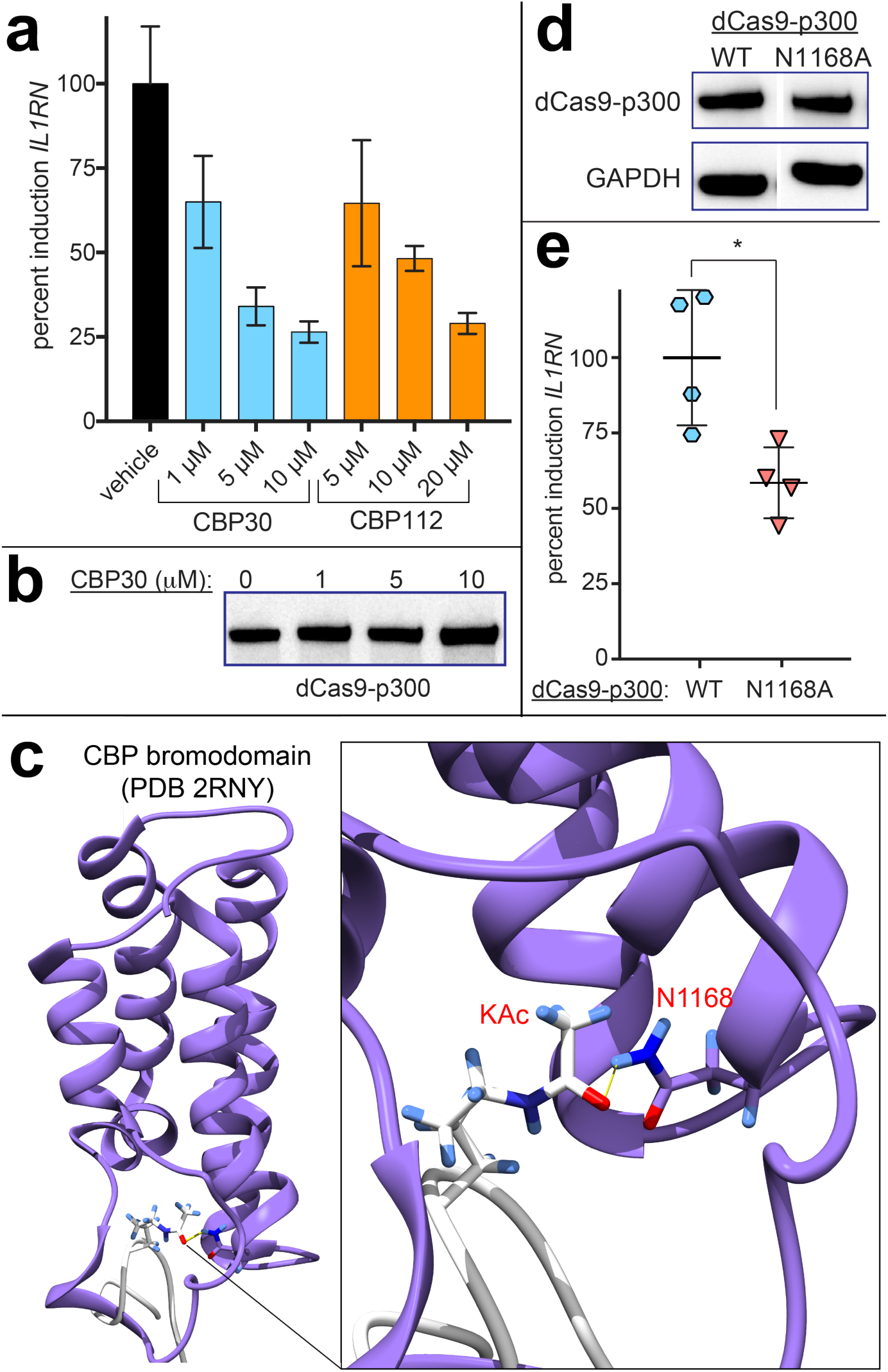
Impact of p300 bromodomain on KAT-dependent gene expression. (a) Dose-dependent inhibition of KAT-dependent gene expression by CBP30 and CBP112. (b) CBP30 does not affect dCas9-p300 overexpression. (c) Crystal structure of CBP bromodomain (PDB 2RNY) highlighting role of N1168 residue (N1132 in full length p300, N1516 in dCas9-p300). in acetyllysine recognition. (d) dCas9-p300 bromodomain mutant (N1168A) mutant is expressed at similar levels to wild-type dCas9-p300. Full gel data is provided in Supporting Information. (e) Mutations known to disrupt the p300 bromodomain-acetyllysine interaction reduce transcriptional activation by dCas9-p300. Significance analyzed by two-tailed Student’s *t* test (* = *P* < 0.05).

In summary, we have described the application of a CRISPR-Cas9 acetyltransferase as a cellular reporter of KAT-dependent transcription. Our studies further establish the scope of this technology, its compatibility with diverse sgRNA architectures, and its application in understanding how small molecule stimuli affect acetylation-dependent gene expression programs. Performing a model screen, we found that dCas9-p300 capably identified ligands interacting directly with p300 (NCGC00496795, CBP30) as well as ligands targeting downstream acetylation-dependent protein-protein interactions (JQ1).^19–20, 23^ This latter finding is both intuitive and striking, as it implies dCas9-p300 establishes an acetylation-dependent “two-hybrid-like” interaction at specific regulatory elements between endogenous cellular components (histones/bromodomains). Based on these results, we anticipate dCas9-p300 should be immediately useful for KAT and BRD inhibitor validation, and potentially amenable to high-throughput discovery applications upon optimization of assay stability and reporter readout.^33^ In addition to identifying new inhibitors, such studies may also provide new insights into KAT-dependent transcription. For example, our results indicate the p300 bromodomain is required for maximal KAT-dependent gene activation at the *IL1RN* locus. Further mutational analysis of dCas9-p300 may aid in the identification of novel determinants of cellular p300 activity. Finally, our studies have implications for small molecule control of dCas9 gene activation. Specifically, we find that dCas9-p300 is inherently ligandable relative to the SAM gene activation system, and also utilizes distinct downstream cofactors (i.e. BRD4). In the future this knowledge could facilitate the development of chemical genetic methods in which tailored inhibitors are used to orthogonally control multiple gene expression programs,^34–35^ and provide an additional strategy for chemical control of dCas9 function.^36–39^ Overall, our studies highlight dCas9-p300 as a powerful tool for studying gene expression mechanisms in which acetylation plays a causal role, and provide a foundation for its future applications in inhibitor validation and biological discovery.

## Supporting Information

Figures S1-S5, Supplementary Tables S1-S4, and supporting materials and methods. This material is available free of charge via the Internet at http://pubs.acs.org.

## Acknowledgements

The authors thank Abigail Thorpe (Chemical Biology Laboratory, NCI) for providing comments on the manuscript and Dr. Steve Kales (NCATS) and Dr. Dominic Esponsito (Protein Expression Laboratory, Leidos Biomedical Research) for helpful discussions. This work was supported by the Intramural Research Program of the NIH, National Cancer Institute, Center for Cancer Research (ZIA BC011488-04) and National Center for Advancing Translational Sciences, National Institutes of Health. In addition, this project has been funded in whole or in part with Federal funds from the National Cancer Institute, National Institutes of Health, under contract number HHSN261200800001E

